# Computationally Enhanced Quantitative Phase Microscopy Reveals Autonomous Oscillations in Mammalian Cell Growth

**DOI:** 10.1101/631119

**Authors:** Xili Liu, Seungeun Oh, Leonid Peshkin, Marc W. Kirschner

**Author notes:** Marc W. Kirschner **Email:**. **Author Contributions** X.L., S.O., and M.W.K. designed research; X.L. performed research; X.L. and S.O. developed image processing algorithms; X.L. developed custom software and analyzed data; X.L., S.O., and L.P. performed periodicity analysis; X.L., S.O., L.P., and M.W.K. wrote the paper.

## Abstract

The fine balance of growth and division is a fundamental property of the physiology of cells and one of the least understood. Its study has been thwarted by difficulties in the accurate measurement of cell size and the even greater challenges of measuring growth of a single-cell over time. We address these limitations by demonstrating a new computationally enhanced methodology for Quantitative Phase Microscopy (ceQPM) for adherent cells, using improved image processing algorithms and automated cell tracking software. Accuracy has been improved more than two-fold and this improvement is sufficient to establish the dynamics of cell growth and adherence to simple growth laws. It is also sufficient to reveal unknown features of cell growth previously unmeasurable. With these methodological and analytical improvements, we document a remarkable oscillation in growth rate in several different cell lines, occurring throughout the cell cycle, coupled to cell division or birth, and yet independent of cell cycle progression. We expect that further exploration with this improved tool will provide a better understanding of growth rate regulation in mammalian cells.

**Significance Statement:** It has been a long-standing question in cell growth studies that whether the mass of individual cell grows linearly or exponentially. The two models imply fundamentally distinct mechanisms, and the discrimination of the two requires great measurement accuracy. Here, we develop a new method of computationally enhanced Quantitative Phase Microscopy (ceQPM), which greatly improves the accuracy and throughput of single-cell growth measurement in adherent mammalian cells. The measurements of several cell lines indicate that the growth dynamics of individual cells cannot be explained by either of the simple models but rather present an unanticipated and remarkable oscillatory behavior, suggesting more complex regulation and feedbacks.

## Introduction

Cell growth is a fundamental physiological property of cells. When cells grow and divide, each component of the cell must double. Although this is well understood for DNA, the strict regulation and coordination of the doubling of all other components is largely a mystery. In non-dividing cells the DNA typically does not double. Yet under these conditions there is still close control of level of all other components. Generally, we can say that cell growth, whether balanced or not, is regulated. In proliferating cell populations, it is strictly tied to cell division, resulting in the control of cell size (1, 2). Deregulation of cell growth is associated with several diseases(3–6). Because for a long time we lacked accurate measurement tools for size of individual cells, cell growth was measured at the population level, as the incremental change in the mean bulk mass or bulk volume in a synchronous culture(7, 8). However, there have long been doubts about the effects of drug induced synchronization of the nuclear mitotic cycle on the rate of cell mass accumulation (8, 9). Some pharmacological treatments used to induce synchrony drive cells from their physiological state, some generate oversized cells which is known to be a distortion of growth(10). Unsurprisingly, bulk studies led to conflicting conclusions, generating controversy. Several approaches have attempted to circumvent the problem of synchronization of a population of cells by finding ways to extract the average growth rate indirectly in an asynchronous population from measurements at a single time point(11–13). Nonetheless, these approaches invariably make questionable assumptions even while calculating population averages. As bulk measures themselves, they may fail to address the role of individual variation in cell growth. Therefore, despite the experimental challenges, it has become more and more clear that it is critical to find ways to measure accurately the growth of single cells over time and then convert this information to population averages if that is desired.

Despite the obvious advantages of single-cell measurements, there are significant experimental challenges. The individual cell growth rate is the change of a single cell mass per unit of time not at a single point in time but at repeated times in the growth phase between divisions. This measurement must be made in an unperturbed way in live cells. It requires extremely accurate size or mass measurement, as the taking of a time derivative greatly amplifies errors. It raises challenges not only in the accuracy but also in the stability, repeatability, and throughput of the size measurement. Cell growth is a collective description of cell mass or cell volume increase. Although cell mass and volume are generally tightly correlated, it is known that volume change can be transient and decoupled from cell mass at different stages of differentiation or of the cell cycle(10, 14–16). We have therefore focused our analysis on the growth of single-cell mass.

There are relatively few methods that accurately measure single-cell mass over time in living cells. The Suspension Microchannel Resonator (SMR) is almost certainly the most accurate. It can measure the buoyant mass to a precision of 50 femtograms or better(17–19), which for a typical cell could be to an error of less than 0.1%. But its application is limited to cells in suspension. It cannot, at present, be used for cell size measurements of adherent cells over a long-time course. Furthermore, in its present form, the SMR only allows for the tracking of a few cells through an entire cell cycle(19, 20) or many cells for a short period of time (21), but not for the most informative tracking of many cells for long periods of time. There are other simpler measurements based on correlations, such as the use of the nuclear area(22) or assaying a constitutively expressed fluorescent protein(23), as proxies for cell mass. However, these proxies are not fully verified and almost certainly are not quantitative enough to make the kinds of critical growth rate measurements that are needed if we wish to reveal small temporal variation in individual cells. In general, correlations between the proxies and cell mass are noisy, reducing our confidence in growth rate measurements. Furthermore, for these methods strict proportionality with mass may not hold for all cell types, at different cell cycle stages, or across the full range of the cell mass distribution. For the reasons above, Quantitative Phase Microscopy (QPM) has emerged as the method of choice for accurate dry mass measurements of attached cells down to a precision of less than ten picograms (note it is still more than 100 times less sensitive than the SMR) (17, 24–26). Furthermore, QPM is more readily available, as subtypes of the QPM technique like the Quadriwave Lateral Shearing Interferometry (QWLSI)(27), Spatial light interference microscopy (SLIM)(28), Ptychography(29), and the Digital Holographic Microscopy (DHM)(30) have been commercialized. An especially attractive form of QPM is the QWLSI, which builds the wavefront sensor around an ordinary Charge-Coupled Device (CCD) and can be easily installed onto a conventional microscope. It can be used with the white-light halogen lamp(27), is compatible with fluorescence detection(31), and has the potential for the high throughput and longitudinal applications. However, in our experience, despite the attractiveness of QPM, the best accuracy can only be achieved with the most optimized yet very restrictive sample configuration and experimental setup. We found that the existing solutions did not provide the stability and robustness required for long–term large-scale growth rate studies to resolve some of the most pressing issues in the field, such as the debate about the linear or exponential growth in single cells(25). Moreover, single-cell growth trajectories usually are complex, noisy, and have large fluctuations(19, 32–34). One needs to evaluate all the sources of error carefully to distinguish spurious fluctuations from meaningful regulation. Finally, both an automated image processing and cell tracking pipeline are required to facilitate the high throughput needed to establish the reproducibility of the measurements.

We report here the development of computationally enhanced reference subtraction and image processing methods for QPM (ceQPM), which improve the accuracy of single-cell dry mass and growth measurements. Specifically, we developed a method to generate a reference phase image to remove the phase retardation unrelated to the sample, improved the algorithm of background leveling, and developed the software for automated image processing, cell segmentation, and cell tracking, all of which enable large-scale longitudinal applications. Using ceQPM, we have carefully quantified the precision of the dry mass and growth rate measurements and successfully monitored the growth rate in thousands of cells in each experiment. The results are sensitive enough to reveal a new feature of cell growth, an unexpected autonomous growth rate oscillation coupled to the cell division cycle.

## Results

### Generating a reference phase image

Quantitative phase microscopy measures the wavefront retardation induced by a sample. It is quantified as the Optical Path Difference (OPD) relative to a reference wavefront(35). However, the OPD of the optical system is often larger than that induced by the sample. Thus subtracting the reference OPD is the most critical step of *quantitative* phase image processing. The reference OPD is generally measured in a cell-free region or from a blank sample. However, this approach is tedious and can be inaccurate in time-lapse imaging because of temporal variation in the system OPD. Here we show the reconstruction of the reference phase image in a more robust manner, which also decreases the noise in the measurement.

When the light crosses the cell area, its phase shifts due to the refractive index difference between the cell and the medium (Fig. 1). Materials in solution maintain a very strict linear relationship between refractive index and concentration. The slope of that relationship is the specific refractive index increment. For proteins, lipids, carbohydrates, and nucleic acids, the specific refractive index increment, α, falls within a very narrow range, with an average of 0.18 *μm*^3^/*pg*(36). The OPD for an entire cell is equal to 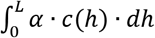, where *c* is the local cell mass density and *L* is the cell thickness. Thus, the total cell dry mas can be measured as

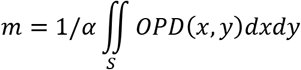

where S is the cell area.

**Figure 1.**
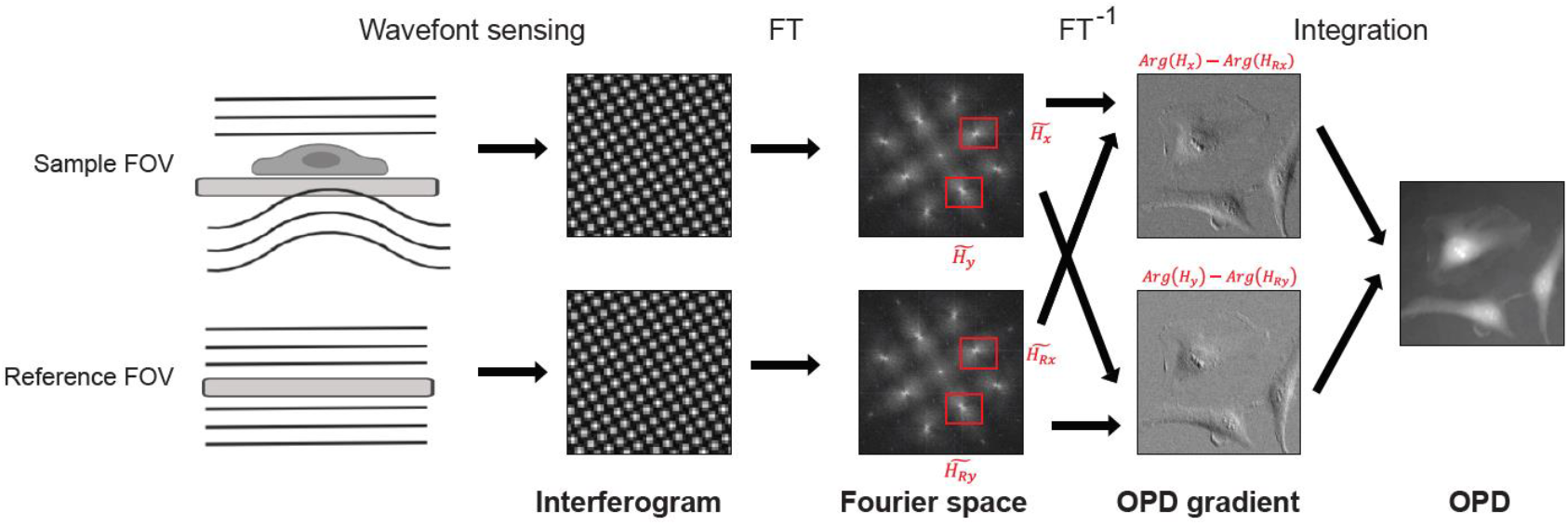
The principle of QWLSI, showing how a reference wavefront is applied to generate the final OPD of the cells.

We used a SID4BIO camera (Phasics S.A., France) to measure the OPD. The camera is based on QWLSI and optimized for biological applications. It uses a Modified Hartmann Mask (MHM) to generate four tilted replicas of the wavefront, which form the interferogram on the CCD sensor. A pair of the first-order harmonics 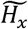 and 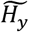 in the Fourier space carries the information for the spatial gradient of OPD in x and y directions (Fig. 1). Thus the OPD is calculated as the 2D integration of the gradients through the Fourier Shift Theorem(27, 37, 38). The resultant OPD contains the phase-shift induced by the cell and an additional phase-shift due to the aberration of the optical system. A reference wavefront is required to remove the phase-shift from the optical system. Knowing the grating period *p* and the distance *z* between the MHM and the CCD sensor, we have

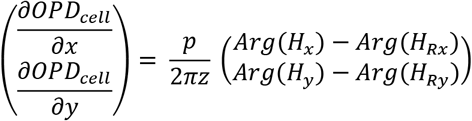

where *H_x_* and *H_y_* are the inverse Fourier transformed images of the Fourier harmonics of the derivatives along x and y of the sample phase image, while *H_Rx_* and *H_Rx_* are the corresponding images of the reference (Fig. 1).

A blank Field of View (FOV) near the sample FOV or an FOV of the same light path on a blank sample is generally used as the reference. However, making a designated blank area in the sample may not always be feasible, and it is tedious and slow to do this manually in large scale screening. We have instead contrived a way to synthesize the reference wavefront. When the confluence of the cells is less than 50%, most of the area of the FOV is blank. Thus we use the median (and not the mean!) of the sample FOVs as the reference wavefront. As *H_x_* and *H_y_* are in complex number form, we calculate their median by calculating the median of the real part and the median of the imaginary part separately.

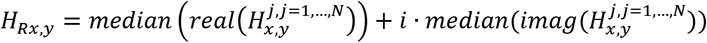

where *N* is the number of sample FOVs. We usually use *N* larger than 16. When the cell confluence is low and all the FOVs share the similar light path (e.g., FOVs on a slide or near the center of a 35 mm or larger dish (Fig. S1C-D)), this method averages out the noise in the OPD measurement and thus performs better than a single reference image of a blank FOV or a blank sample (Fig. S1A). Both the cell confluence and the similarity in the light path affect the goodness of the synthetic reference (Fig. S1B, C, and D). For the best performance of the method, we usually seed cells at lower than 30% confluency and scan within the central 10% area of a circular dish or well. It is worth mentioning that the rationale for generating the synthetic reference is not limited to QWLSI. The reference light path at any time during a time course can be retrieved through a similar method for other types of QPM. This is especially helpful for the QPM systems more subject to reference path distortion due to the use of coherent light sources or long reference arms.

We developed Graphical User Interfaces (GUI) as well as scripts to generate position matrix of desired pattern, make synthetic reference, and evaluate the performance of the reference image before applying it to the whole data set. All of these make the high throughput QPM measurement much more convenient and improve the reproducibility and the accuracy of the measurement.

### Background leveling corrections

As shown in Fig. 2A, there is still residual background after compensating for optical system aberration by subtracting the reference image. The residual shape of this background could be due to cover glass thickness variation, vibration, etc. Background leveling is crucial for accurate dry mass quantification. The conventional methods of polynomial fitting(25) or Zernike polynomial fitting(39) capture the low-frequency background but miss the regional fluctuations (Fig. 2B). We developed a new method for subtracting both the low and high-frequency background, thus significantly improving the precision of the dry mass measurement.

**Figure 2.**
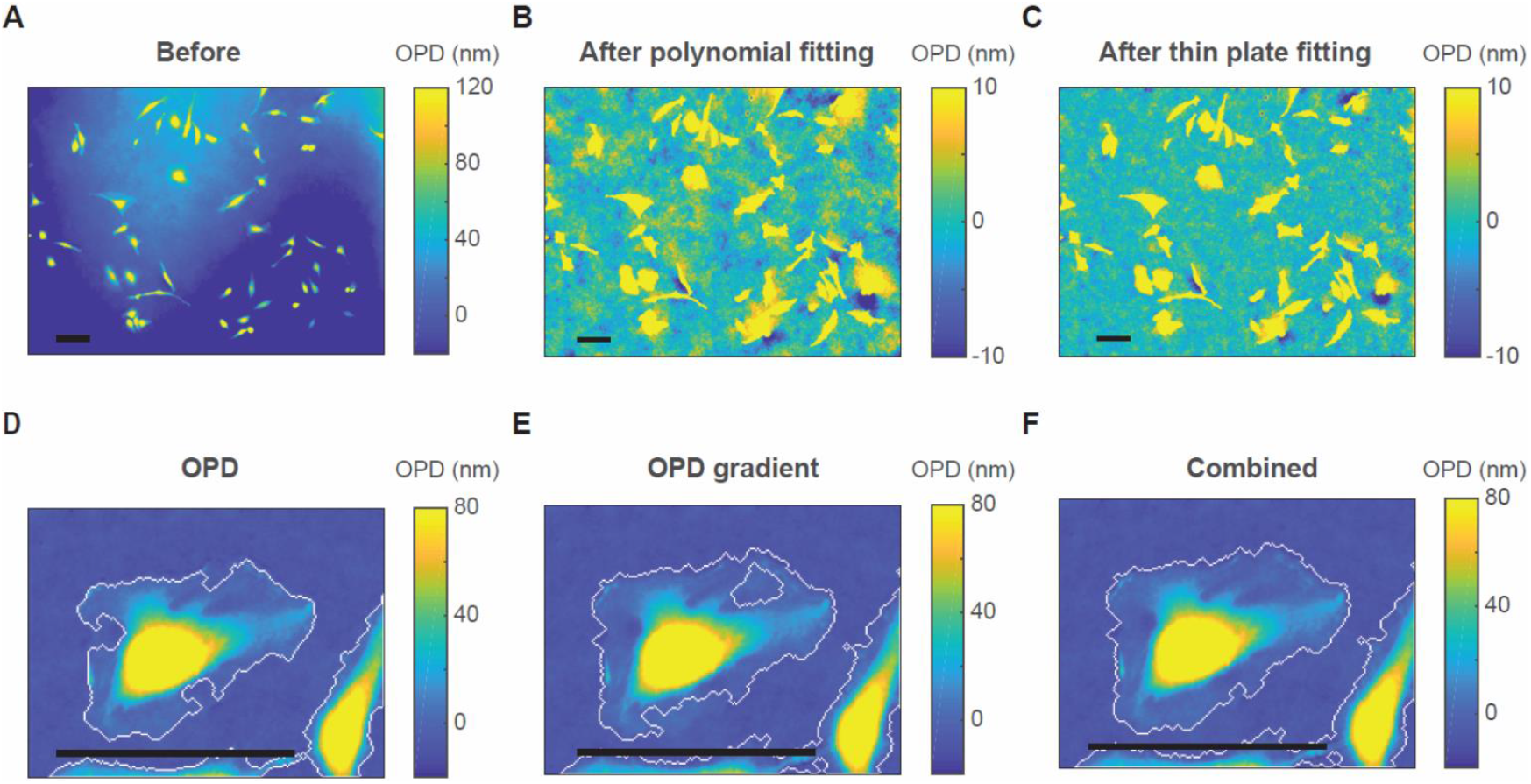
Background leveling. (A) The OPD image before background leveling. (B) The OPD image after subtracting the background fitted by a 2D polynomial (n = 8). (C) The OPD image after subtracting the background fitted by the thin plate spline. (D, E) Cell boundaries determined by a threshold on the OPD images (D) or the gradient magnitude of the OPD image (E). (F) The combination of the boundaries on (D) and (E). (A-C) are from an FOV under a 10X objective lens. (D-F) are from an FOV under 40X objective lens. Scale bars indicate 100 µm.

We first isolate the objects from the background by “top-hat filtering”. A disk-size smaller than most of the cells is chosen as the structuring element to clean up the fluctuations whose scale is comparable to or larger than the cell. The resultant image cannot be directly used for quantification because it subtracts excessive background from the cells, and the mean of the background level varies with each image. We use it only to generate the background mask: we separate the image into the cell and the cell-free areas by combining the filtered OPD image and its gradient magnitude to define the boundary of the cells. Thresholding OPD or OPD gradient alone may leave out part of the cell (Fig. 2D and E), but the combination of the two detects the cell boundaries much more accurately (Fig. 2F). Note that we intentionally do not fill the holes in either thresholding mask, as other QPM segmentation methods recommended(25, 39), because this process may also fill the blank area within a cell cluster, which is critical for the precise fitting of the cell-dense area. Lastly, we create the background image by fitting the cell-free area of the *original* image with a thin plate spline method(40). A mesh grid binning is used for fast computation. The thin plate fitting is parameter-free and can capture both large and small curvatures. Fig. 2C shows that our method generates a cleaner background than conventional methods.

### The precision of dry mass measurements

The dry mass measurement error contains all the variation in the OPD measurement, the background subtraction, and the cell segmentation. Among those factors, the background subtraction has the largest effect, as the unevenness in the background affects the cell boundaries. We quantified the precision of the dry mass measurement by measuring same fixed cells multiple times at different positions. The result is summarized in Table 1. Our background subtraction algorithm significantly improved the precision of the dry mass measurement. The combined temporal and spatial error is reduced at both high (40X) and low (10X) magnifications when compared to a previous study using a similar setup but different data processing algorithms (Table 4 in ref. 25). Remarkably, the error at 10X is reduced by more than two-fold from 4.37% in Aknoun et al.(25) to 1.97% in our study. As the magnification decreases from 20X to 10X, the FOV is four-time larger, while the measurement error increases only 1.15 folds. We, therefore, gained acquisition throughput without sacrificing much precision of measurement. For this reason, we optimized our data collection to maximize throughput at 10X.

**Table 1.**
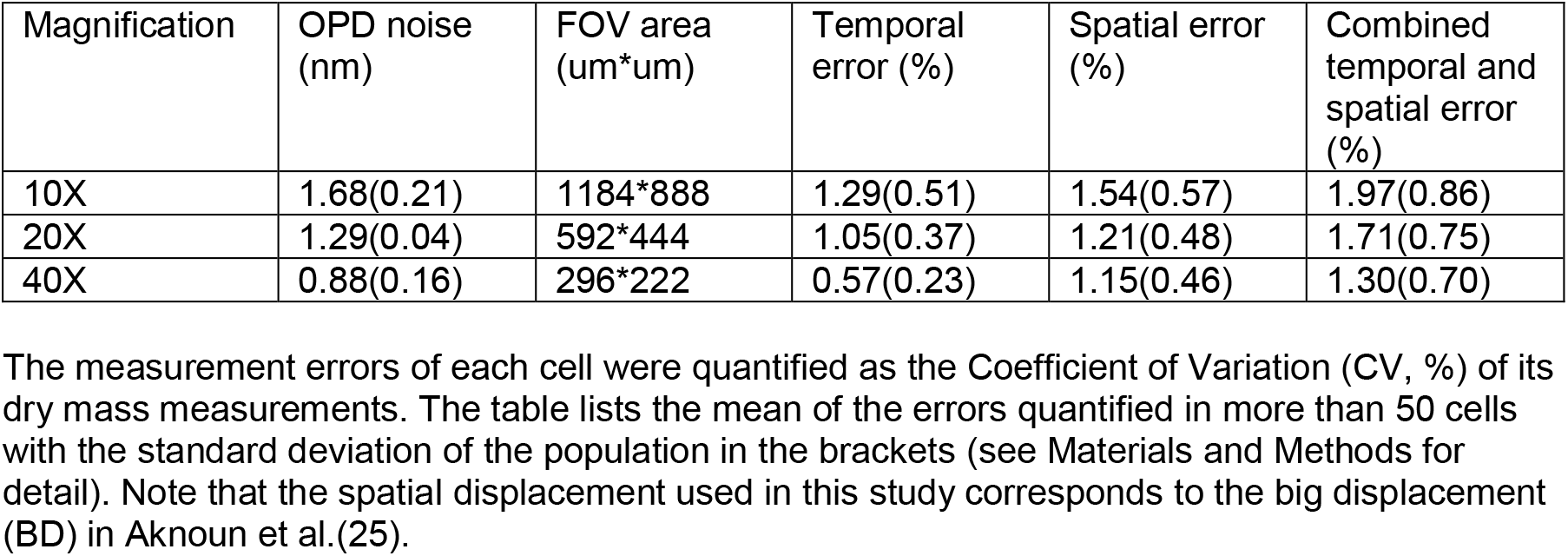
Dry mass measurement precision at different magnifications.

### Cell segmentation and cell tracking

For cell segmentation, the watershed algorithm works the best when a nuclear marker is used as the foreground marker(41). When no nuclear marker is available, we use the local maximum of the cell after top-hat filtering. Because two cells may closely contact each other and form only one local maximum or one cell may have two local maxima, segmentation without any nuclear marker possesses about 5% error depending on cell types. We now find it most useful to combine the OPD image and its gradient magnitude to define the boundary of the cells as discussed in *Background leveling corrections*.

To track cell mass in time automatically, we first identify all the mitotic cells in the time series based on their rounded morphology, by their mass density gradient and area (Fig. 3A). We then trace cells backward. Each track starts from the end of the time series or a mitotic event. No new track is added during the tracing process. We use cell mass, area, and centroid position as the metrics for tracing. We compare a cell *k* on frame *i* with each cell on frame *i* − 1 by the weight function:

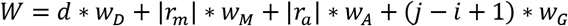

where *d* is the distance between the centroids, *r*_*m*_ is the relative mass difference, *r*_*a*_ is the relative area difference, *j* indicates when the metrics of cell *k* were last updated, and *W*_*D*_, *W*_*M*_, *W*_*A*_, and *W*_*G*_ are the weights of the respective terms. The dry mass measurement is so precise that we can put high confidence in the mass term. The weight parameters for HeLa are summarized in Table S1, as an example. The value of *W* is used to determine the goodness of the match. A good match should have the smallest *W* on the frame and *W*<1. When cell *k* has a good match, its metrics are updated with the newly traced cell. Otherwise, the old metrics are carried on for comparison with the next frame. This method may leave gaps in tracks that can be filled later by a smoothing algorithm but tolerate most of the segmentation error. The track does not terminate or deviate by wrong segmentation of a single frame. A track essentially terminates when it cannot find a good match for more than ten frames (*j* − *i* + 1) * *W*_*G*_ > 1. In the last step, we trace the lineages of the cells. We compare the metrics at the end of each track with all the mitotic cells. If a track ends just before a mitotic event (the time axis is inverted), the centroid position is near the position of the mitotic cell, and the mass is close to half of the mitotic mass, the track is identified as the daughter cell of the mitotic cell. Because newborn cells tend to contact their sisters closely, this will result in problematic segmentation. For that reason, many cells cannot be traced back to the very beginning of birth. For consistency, we designate the division time (the last frame of mitotic rounding) of the mother cell as the birth time of the daughter cells.

**Figure 3.**
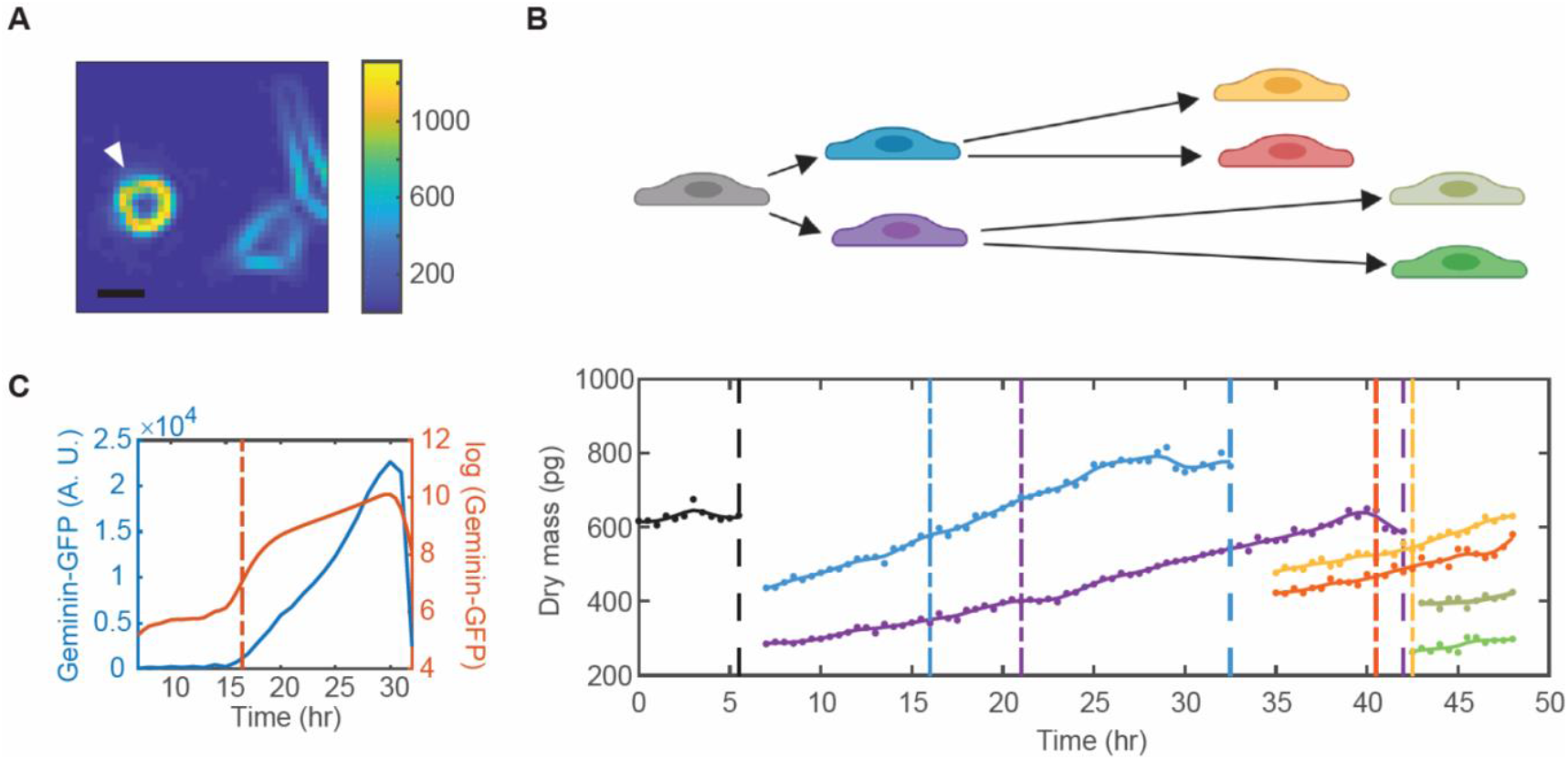
Cell tracking. (A) The gradient magnitude of an OPD image measured at 10X. Scale bar indicates 20 µm. The arrow indicates a mitotic cell. (B) One cell is traced to its granddaughter cells. Each color represents a cell. Solid dots are the raw data of dry mass measurement. Solid lines are the spline line smoothing. Vertical dashed lines indicate the timing of cell divisions. Dash-short dashed lines indicate the timing of G1/S transitions. (C) The intensity of Geminin-GFP measured in one cell (blue) and its logarithm (red). Dash-short dashed line indicates the steepest slope of the log(Geminin-GFP) accumulation curve, which is defined as the time of the G1/S transition.

Fig. 3B shows an example of a cell traced to its grand-daughter cells. For each cell, the G1/S transition is determined by the steepest slope of log(Geminin-GFP) accumulation curve (Fig. 3C). Using the methods described above, we were able to successfully trace the cells in a completely automated fashion without manual supervision or correction. The fraction of mistraced cells identified by manual counterchecking is less than 2%. Movie S1 shows an imaging area of 9.47 mm × 7.99 mm on one well of the 6-well plate monitored for 48 hours under 10X objective. A total number of 2983 cells were traced in the area. Six imaging areas of such can be measured within 30 minutes. The limit of the measurement throughput is the speed at which we can move the stage without perturbing the optical stability of the culture medium surface (see Materials and Methods for specification) and the rotation speed of the filter turret.

### Evidence for growth rate oscillations during cell growth

It has been a long standing question whether the growth of individual proliferating cell can be described as linear or exponential (7). It is surprisingly difficult to distinguish the two models, because cell size just doubles in one proliferation cycle rendering the maximum difference between the two models only 5.63%(42, 43). Since our measurement error is lower than 2%, it allows us to address this question in adherent mammalian cells, when monitored throughout the cell cycle. To assure the highest measurement accuracy, we cleaned up the individual cell growth trajectories by eliminating any data points where the cell was in contact with another cell, as the cell-to-cell contact usually causes erroneous segmentation of more than 2% dry mass. Furthermore, we eliminated rounded cells, as their dry mass error is also larger due to the dramatic change of height and the problem of phase unwrapping. In most trajectories, the first 1.5 hours after birth and the last 0.5 hours before division were removed during this cleaning process. As a result, the cleaning process discarded most of the cell trajectories, and less than 10% were used in the following analysis. We collected 340 full-cell-cycle trajectories of HeLa cells from three replicative experiments and pooled all of the trajectories together to investigate which model explains the growth dynamics better. We compared the goodness of fit by the small sample Akaike Information Criterion (AICc); the better fit possessed the smaller AICc(44) (Supplementary Information). The exponential model fitted much better than the linear model in the mean trajectory of the whole population (Fig. 4A, ∆AICc = 102.97), which was consistent with the positive correlation between growth rate and cell size found previously(33). Yet, neither model fits every cell. The exponential model fits better in 68.6% of the population, whereas the linear model fits better for the rest (Fig. 4B). The ratio was similar in another adherent cell line, RPE1 cells, where 61.5% cells got fit better by the exponential model (Fig. S2A).

**Figure 4.**
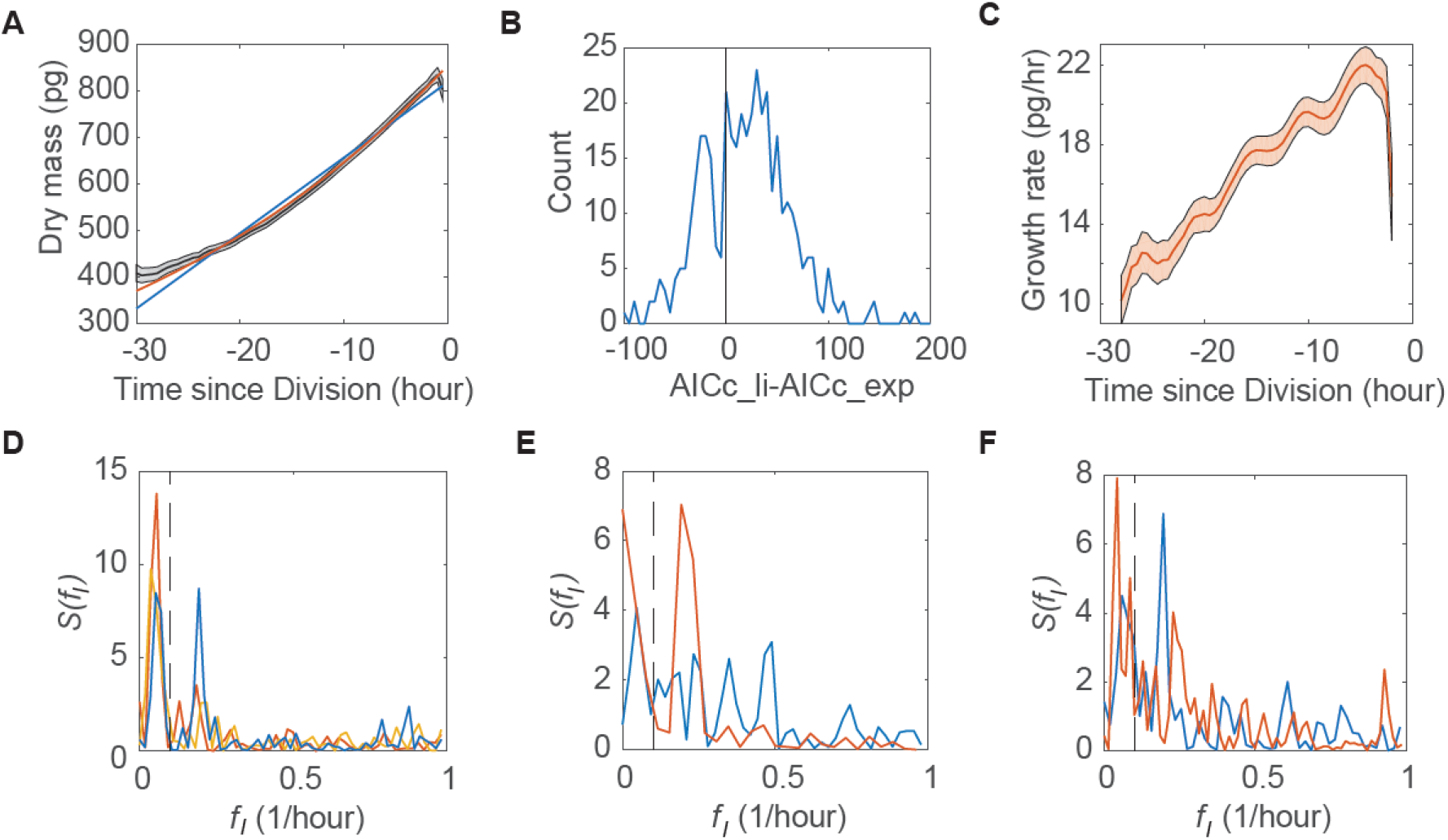
Detection of the growth rate oscillations in adherent cells. (A) The mean trajectory (black) of the smoothed dry mass trajectories of HeLa cells aligned to cell division time, fitted by the linear (blue) or exponential (red) growth model. Data were collected from three independent replicative experiments, n = 340. The gray shaded regions indicate the 95% confidence intervals of the mean. (B) The histogram of the difference between the AICc of the linear (AICc_li) and exponential (AICc_exp) fit in HeLa cells. Better fits have the smaller AICc. The black vertical line indicate the difference equal to zero. (C) The mean trajectory of the smoothed growth rate trajectories of HeLa cells aligned to cell division time. The red shaded regions indicate the 95% confidence intervals of the mean. n = 340. (D) The periodogram of the mean trajectories of the detrended dry mass trajectories of HeLa cells aligned to cell division (blue), birth (red), and G1/S (yellow), respectively. n = 340. (E) The periodogram of the mean trajectories of the detrended dry mass trajectories of HeLa cells under thymidine treatment aligned to their chronological time (blue) or the first peak after G1/S transition (red). n = 51. (F) The periodogram of the mean trajectories of the detrended dry mass trajectories of HeLa cells under control condition (blue) or rapamycin treatment (red) aligned to cell division time. n = 188. The periodograms in (D-F) were estimated by the robust detection method as described in Ahdesmäki et al. (47). The black dashed lines indicate *f_min_*=0.1/hour used in the g’-statistic test.

Growth dynamics were more complex than any of these simplified models. We found the residual of the fit in some single-cell trajectories seemed to be oscillatory, which was particularly intriguing (Fig. S2B). The oscillatory behavior is the most apparent in the mean of the growth rate trajectories aligned to cell division time (Fig. 4C). In our initial analysis, we smoothed the mass trajectories and took their time derivative by fitting the linear slope in a short sliding time window. These manipulations of the data should not change the overall shape, exponential or linear, but the smoothing and derivative taking could produce artifactual periodicity (Supplementary Information).

To investigate the unexpected growth rate oscillation, we therefore went back to the original unsmoothed raw dry mass data. Proof for periodicity in noisy observations has been thoroughly considered in astronomy as well as in biology(45, 46). It’s important to formulate the problem of the existence of periodicity as a statistical question and test it against a null hypothesis model of random fluctuation, which is what we have done in this study. Specifically, we used the robust detection method developed by Ahdesmäki et al. (47). It derives the “robust” periodogram from the correlogram spectral estimator and tests the significance of the maximum peak against the null hypothesis of randomly permutated data. This method has special advantages as it is insensitive to outliers, applies to short time series, and does not require assumptions on the form of noise. Similar to Fisher’s g-test (48), this method defines the g-statistic as

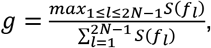

where *S*(*f*_*l*_) is the spectral estimation at the frequency *f*_*l*_, with 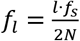, *l* = 0,1,…,(2*N* − 1)/2, *N* is the length of the trajectory, and *f*_*s*_ is the data acquisition frequency. *f*_*c*_ denotes the frequency at the maximum of *S*(*f*_*l*_). If the trajectory is determined to be oscillatory, *f*_*c*_ equals the oscillation frequency. The p-Value of the observed g-statistic was estimated by 5000 randomly permutated trajectories. It allowed us not only to investigate the dominant 5-hour period oscillation which we found in the average growth rate trajectory, but also to discover oscillations of any frequency and amplitude if they were more significant than noise. We first detrended individual dry mass trajectories by the second-order polynomial, which fits better than either of the exponential or linear model in 63.9% cells (Fig. S2D). Then we aligned those trajectories to cell division and averaged them (Fig. S2E). The periodogram of the mean trajectory presented two distinct peaks (Fig. 4D). The peak at 0.053/hour (19.0-hour period) could be due to the imperfect detrending or actual growth rate slowdown in the middle of cell cycle. As the period of this peak is close to the average length of the cell cycle (26.2 hours for HeLa cells), we designated it the “cell cycle” peak. The other peak at 0.193/hour (5.2-hour period) corresponds to the periodic bumps in the mean growth rate trajectory in Fig. 4C, which we designated as the “sub-cell cycle” peak. The “sub-cell cycle” peak is more dominant, its p-Value is 0.0014, meaning that the about 5-hour periodicity is highly significant.

However, the “cell cycle” peak affects the significance of the “sub-cell cycle” peak. Especially when the “cell cycle” peak becomes the maximum peak in the periodogram, the robust detection method tests its significance rather than the “sub-cell cycle” peak. Since the “cell cycle” peak could be due to the imperfect detrending and we were more interested in the “sub-cell cycle” peak, we revised the question from whether there is a significant peak in the periodogram to whether there is a significant peak beyond the frequency *f*_*min*_, where the choice of *f*_*min*_ is chosen ad hoc to be higher than the “cell cycle” peak but lower than the “sub-cell cycle” peak. The period of the “cell cycle” peak can be longer or shorter than the cell cycle length, but is always longer than 10 hours, we used *f*_*min*_ = 0.1/*hour* in most cases unless specified otherwise. As the robust detection method is insensitive to the choice of the *l* series, we can answer the new question within the same framework by modifying the g-statistic to

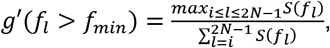

where *i* is the smallest *l* of *f*_*l*_ > *f*_*min*_. We validated the g’-statistic by investigating the False Discovery Rate (FDR) in random trajectories with Gaussian noise as well as mean trajectories of permutated single-cell trajectories. The FDR in both datasets was close to 5% when the p-Value was set to 0.05 (Supplementary Information). According to the g’-statistic, the p-Value of the “sub-cell cycle” peak is less than 0.0002 (i.e. none of the 5000 permutated trajectories has a larger g’), showing the extraordinary goodness of the periodicity. In the following analysis, we alternatively used the g- or g’-statistic depending on the existence of the “cell cycle” peak, whose p-Value were defined as *p1* and *p2*, respectively.

Periodic variation could be induced by fluctuations in instruments, particularly if they coincided with built-in variation in hardware, like temperature, light, line voltage, etc. We used the fixed-cell data monitored for a long time to investigate the instrumental fluctuation. Unlike the live-cell data, the fixed-cell data is not oscillatory (*p1*=0.4334) (Fig. S2E, F). As proposed by Ahdesmäki et al. (47), we could use

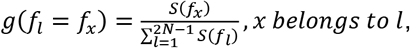

to test the significance at a specific frequency, *f*_*x*_. The p-Value of *g*(*f*_*l*_ = *f*_*x*_) was defined as *p3*. According to this test, there is no significant peak near *f* = 0.2/*hour* (*p3*=0.9999) in the periodogram of fixed cells. Hence, we concluded the ~5-hour period in HeLa cells was not due to the instrumental fluctuation.

We next tried to identify oscillation in individual cells. The g-statistic found 40 oscillatory cells out of 564 with *p1* smaller than 0.05 in the fixed-cell data, which occupied 7.1% of the population, whereas it found 106 oscillatory cells out of 340 in the live-cell data, which occupied 31.2% of the population (Fig. S3A, B). Since the oscillations of frequency smaller than 0.1/hour could be due to imperfect detrending, we focused on the 62 oscillatory cells out of 340 of frequency larger than 0.1/hour, whose percentage (18.2%) is still significantly larger than the 7.1% in fixed cells (p-Value = 7×10^−7^ by Fisher’s exact test). Their average oscillation frequency was 0.195/hour, which was very close to the 0.193/hour oscillation frequency of the mean trajectory of the whole population. The higher noise level of single cell trajectory may have concealed the oscillation in the 68.8% of cells that did not meet p<0.05 criteria. To probe whether these cells had oscillation, we removed the 106 oscillatory cells from the dataset and took the mean trajectory of the remaining cells. g’-statistic test of this mean trajectory showed oscillatory (*p2*=0.0018) (Fig. S3C), suggesting that the growth rate oscillation in Fig. 4D was not caused by a subpopulation of cells but rather existed in the whole population. It is worthy noting that, the oscillation amplitude of the mean trajectory is less than 0.5% of the average cell mass (3 pg vs 600 pg), which is much smaller than the 2% measurement error of ceQPM in single cell trajectories. When the measurement error was larger than the oscillation amplitude in single cells, the characteristic peak of the oscillation in the periodogram might be masked by random noises. However, if all cells were oscillating in similar frequency and properly aligned by their oscillation phase, the random noise in the mean trajectory would be averaged out, leaving the oscillation peak distinct in the periodogram. This may explain why we didn’t detect oscillation at the single-cell level (but did in their mean trajectory) of the 68.8% cells. If it were the case, one would expect reducing the number of the constituent trajectories or aligning them to a different time could perturb the oscillatory behavior in the mean trajectory. Indeed, we examined the mean trajectory of a single experiment which happened to provide half of the data (170 cells). When the trajectories were aligned to their division time, in its periodogram, the maximum peak of *f_l_*>0.1/hour was at 0.196/hour, consistent with the average of three experiments. Due to the smaller size of data, there were more random peaks and *p2* increased to 0.0840 (Fig. S3D). When we aligned those trajectories to their chronological time rather than cell division, the ~0.2/hour peak disappeared (Fig. S3E). Instead, a distinct peak showed up at 0.983/hour (*p1*<0.0002). During the experiment, we acquired the phase images every 30 minutes and the fluorescent images every 1 hour. The microscope spent longer time to scan all the positions with both channels than just one channel, thus the time interval in the trajectories was not even. However, we assumed it to be even in the data analysis, leading to the 1-hour artificial oscillation. Our method was sensitive enough to pick up this subtle oscillation but indicated no significant peak around the 0.2/hour frequency (*p3*=0.9999). This result further confirmed that the ~5-hour-period oscillation in live cells was not introduced by environmental cues but was intrinsic to growth rate regulation, and its phase was tightly coupled to cell division.

We next examined the oscillation phase relative to cell cycle events other than cell division. When we aligned all the 340 trajectories of the three experiments to cell birth, the ~0.2/hour peak in the periodogram was preserved (*f*_c_=0.185/hour) but became less significant (*p2*=0.0410), whereas when we aligned them to G1/S transition, the mean trajectory was not oscillatory any more (*p2*=0.2849) (Fig. 4D). We found similar results in RPE1 cells (Fig. S3F). When the detrended cell mass trajectories were aligned to cell division time, the mean trajectory was almost oscillatory (*p2*=0.0794) with two distinct peaks of *f_l_*>0.1/hour. The slightly higher peak was at *f*_c_=0.324/hour, while the other peak was at 0.216/hour, which was close to the ~0.2/hour peak in HeLa cells. When the trajectories were aligned to cell birth or G1/S, the mean trajectory was not oscillatory with *p2* of 0.7856 or 0.3647, respectively. Note that the cell number in the RPE1 dataset is smaller than in the HeLa dataset, and the two distinct peaks of similar height compromised the significance of periodicity in the test used currently, which was designed to detect single oscillation frequency. Nonetheless, the main conclusion was the same as HeLa cells that the oscillation in the mean trajectory was much more pronounced when the cells were aligned to division than to birth or G1/S, suggesting a coupling between the growth rate oscillation and the cell division cycle. We further investigated Hela cells under thymidine treatment. We aligned all the trajectories to the first peak after G1/S and found that the oscillation proceeded after the cells entered into S phase and arrested (*p2*<0.0002) (Fig. 4E). It implies that the growth rate oscillation is autonomous and independent of cell cycle progression. Here, we point out that cell growth continued when cell cycle was arrested at S-phase (Fig. S3G). As growth rate is the net difference of protein synthesis and degradation, drugs inhibiting protein synthesis rate may also perturb the oscillation. Indeed, under treatment with rapamycin, the mean trajectory was less oscillatory compared to the normal growth condition of the same cell number (*p2*=0.2487 vs *p2*=0.0016) with the *f*_c_ shifted from 0.1961/hour to 0.2286/hour (Fig. 4F).

All of these experiments above were done with adherent cells measured by ceQPM. We then decided to investigate if we could see any oscillations in growth rate of suspension cells measured by the much more accurate SMR. We are grateful for the sharing of a large set of data of a mouse lymphoblastoid line, L1210, measured by Mu et. al.(49). Similar to the QPM data, we first detrended the individual buoyant mass trajectories by the third-order polynomial, which fitted better than either of the linear or exponential growth models in all the cells (Supplementary Information). Since the SMR data were measured at such high accuracy and fine time resolution, we were able to reveal the periodicity in single cells. The robust detection method found all of the 63 cells we analyzed were oscillatory with *p1* less than 0.0002. Each cell only had one or two outstanding peaks in its periodogram (data not shown). Then we fitted individual trajectories with the generic cosine function,

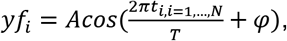

where *t*_*i*,*i*=1,…,*N*_ is the time series, *N* is trajectory length, *A* is oscillation amplitude, *T* is oscillation period, and *φ* is the phase at *t*_*i*_ = 0 (Fig. 5A-F). We evaluated the goodness of fit by the adjusted Residual Sum of Square (adj_RSS),

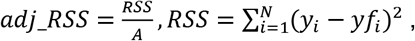

where *y*_*i*_ is the observation, *yf*_*i*_ is the fitted result. Fig. 5G shows the distribution of *adj_RSS* of the 63 cells. We arbitrarily chose a cutoff threshold at *adj_RSS*=50 and only investigated the fitted results of the 56 out of 63 cells below the threshold. Note that the cells with *adj_RSS* above 50 were also oscillatory but with bigger variation (Fig. 5F). We found the average period was 3.6 hours (Fig. 5H) and the average amplitude was 0.11 pg (~0.2% of the averaged cell mass) (Fig. 5I). Unlike the adherent cells, in L1210, the oscillation phase was tightly coupled to cell birth but not so related to G1/S transition or cell division (Fig. 5J). As the SMR data traced the cell lineage for several generations, we also investigated the mother-daughter correlation of the oscillation properties (Fig. 5K-N). We found positive correlations in oscillation amplitude and period among the mother and daughter cells (Fig. L, M). However, the daughter cell does not inherit the phase from its mother, meaning the oscillation phase was reset at cell birth per each generation (Fig. K, N). We further calculated the Pearson’s correlation between all the measured variables. Fig. 5O summarized all the correlations of p-Value smaller than 0.05. We found that period was strongly correlated with cell cycle length, and hence negatively correlated with cell size. Amplitude had a positive correlation with cell cycle length and period. The phase at birth was positively correlated with division mass.

**Figure 5.**
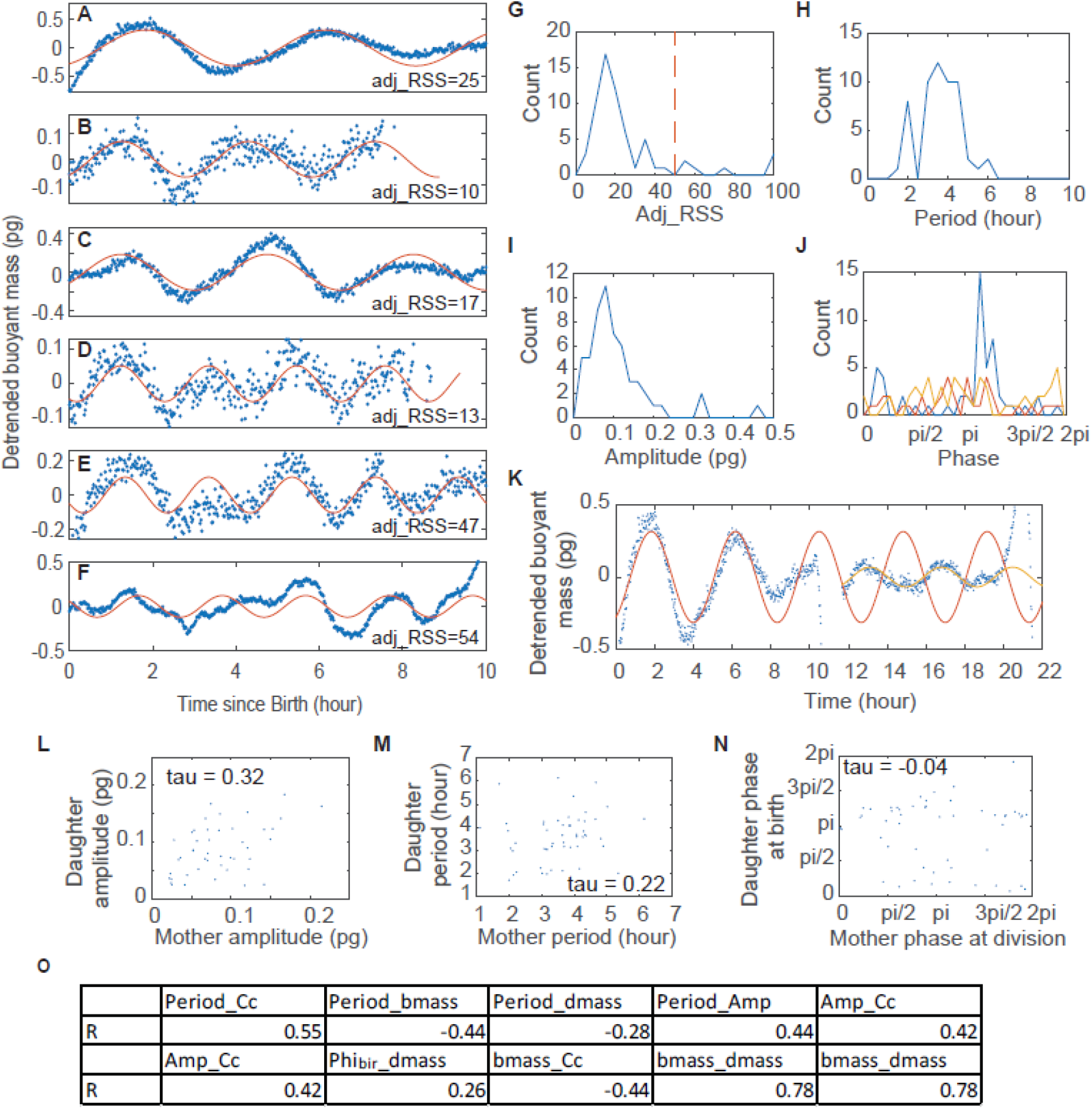
Detection of growth rate oscillations in individual L1210 lymphoblast cells. (A-F) Examples of randomly selected detrended buoyant mass trajectories (blue dots). The solid red lines are the fitted cosine functions. adj_RSS indicates the goodness of fit. (G-J) The distributions of adj_RSS (G), period (H), amplitude (I), and phase at different cell cycle events (J) of the fitted cosine functions. The red dashed line in (G) indicates the arbitrary cutoff threshold; the distributions in (H-J) only include the 56 cells below that threshold. The blue, red and yellow lines in (J) represent the oscillation phase at birth, G1/S, and division, respectively. Note that only 38 cells were measured with the fluorescent FUCCI marker and had identified G1/S timing. (K) The detrended buoyant mass trajectory of an example L1210 cell continued with the trajectory of one of its daughters (blue dot). The red and yellow lines are the fitted cosine functions of the mother and daughter cell, respectively. (L-N) The mother-daughter correlation of the oscillation amplitude (L), period (M), and phase (N). The correlation coefficients were calculated as Kendall’s Tau coefficient to avoid the effect of outliers. (O) The summary table of Pearson’s correlations of p-Value smaller than 0.05. Cc denotes cell cycle length; bmass denotes birth mass; dmass denotes division mass; Amp denotes amplitude; Phi_bir_ denotes the phase at birth.

In summary, we identified the growth rate oscillations in two adherent cell lines, HeLa and RPE1, and one suspension cell line, L1210, which were of comparable period (3.1-5.2 hour) and amplitude (0.2-0.5% of the averaged cell size). The oscillations were measured by the two most accurate but totally distinct methods at different time resolutions, confirming the periodicity is very real and virtually impossible to be caused by experimental or analytical artifacts.

## Discussion

How cell growth and cell division are coordinated has turned out to be a very difficult problem. The difficulty arises primarily from the challenge of making accurate measurements of cell mass over time in single cells. The most commonly used and thoroughly studied mammalian cells are the cells attached to plastic or glass. They have many experimental advantages for some technologies but fail to others. QPM has become the optimal method to measure cell mass in adherent cells(17, 24, 32, 50, 51). Yet its sensitivity was not sufficient for deciding growth models, such as whether for proliferating cells it is linear or exponential. We have devised computational innovations in ceQPM that can significantly improve the measurement accuracy by more than two-fold. ceQPM also improved the stability and repeatability of the measurement. It allowed us to use lower magnified lenses covering larger field and producing more measurements per unit time. The advanced data acquisition throughput provided high statistical power. The automated cell segmentation and tracking algorithms facilitated the processing of large datasets. Both the experimental and analytical systems can be incorporated into wide applications of cell size and cell growth studies. We used this improved method to explore the growth of two adherent cell lines, HeLa and RPE1 cells. Both cell lines are more closely fit to an exponential model. Yet, neither the exponential nor the linear model fits all the cells. Rather the exponential model fits better in about two thirds of the population.

When we looked carefully at the growth curves, we found clear hints of some oscillatory behavior. On deeper analysis there was a 3.1 to 5.2-hour oscillation in growth rate in HeLa and RPE1 cells, which is coupled to cell division time but unrelated to the G1/S transition of the cell cycle. Note that although later pooled and synchronized *in silico*, these cells were measured from *asynchronous* population in different experiments so that the experimental time bears no relationship to real-time and the individual members of the population can be expected to be scrambled with respect to time in each experiment. It rules out the possibility that the periodicities arose from environmental fluctuation (such as temperature or line voltage) or some collective signal among the cells in the well. We also observed a growth rate oscillation of comparable period and amplitude in a suspension cell line, L1210 lymphoblast, measured by SMR. The SMR, having at least 100-times better accuracy and 25-times higher time resolution than the ceQPM measurements, detected the periodicity in almost all the individual cells. At this point, we do not know what drives the oscillations or whether they are of the same underlying mechanisms. The limited pharmacological perturbations provide some hints. The blockage of the nuclear division cycle with thymidine did not arrest the oscillation, suggesting that the oscillation is independent of cell cycle progression. The partial inhibition of growth with rapamycin weakened the periodicity and shortened the period from 5.1 hours to 4.4 hours, suggesting a plausible linkage to protein synthesis, degradation, or metabolic activity through the mTOR signaling pathway. Further study of correlation between the oscillation parameters (amplitude, period, and phase) and cell properties (cell mass, cell cycle length, etc.) could provide a means to a better understanding.

Although the growth rate oscillation is autonomous, its phase is coupled to cell division in HeLa and RPE1 and to birth in L1210 cells, which suggests possible mutual entrainment between the growth rate oscillation and the cell cycle. The growth rate oscillation may serve as a gate to mitosis by controlling the availability of metabolites and cellular energy level. On the other hand, mitosis, as the most dramatic event in the cell cycle, might reset the growth rate oscillation at birth by pausing RNA and protein synthesis or even depleting cellular ATP(20, 52). Investigating the coupling between cell growth and cell cycle oscillations could provide novel understandings of both(53–57). The coordination between the two is critical for cell size regulation and may shed light on the cause of heterogeneous drug response among isogenic cells in cancer therapy.

Usually, an oscillation is caused by negative feedback with a substantial time delay(58). Due to the complexity of biological networks, oscillations may broadly exist. Among them, the periodic protein synthesis has been reported several times in drug-induced synchronous cultures(59–62), and the periodic cell size changes or protein production rate caused by metabolic oscillation was also discovered in individual budding yeast cells(53, 63). However, the idea of endogenous growth rate oscillation has never been widely considered due to the lack of convincing experimental evidence. In this study, we discovered the oscillations in unperturbed cells of two adherent cell lines measured by ceQPM and one suspension cell line measured by SMR, suggesting that periodicity may be a general property of growth dynamics and exist in non-dividing cells. Although subtle, growth rate oscillations may hint at a system for maintaining cell size and growth homeostasis.

## Materials and Methods

### Cell culture

HeLa Geminin-GFP and RPE1 Geminin-GFP cells were generated and single clones were isolated and grown in our laboratory(12). Cells were kept in Dulbecco’s Modification of Eagles Medium (DMEM, ThermoFisher Scientific, 11965), supplemented with 10% Fetal Bovine Serum (FBS, ThermoFisher Scientific, 16000044), 1% penicillin/streptomycin (10000 U/mL, ThermoFisher Scientific, 15140122), 10 mM sodium pyruvate(100 mM, ThermoFisher Scientific, 11360070), and 25mM HEPES (1 M, ThermoFisher Scientific, 15630080), and incubated at 37°C with 5% CO_2_. Rapamycin used to inhibit mTOR activity was purchased from LC Laboratories (R-5000). Thymidine to arrest cell cycle was purchased from Sigma-Aldrich (T1895).

### Microscope setup

The SID4BIO camera (Phasics, France) was integrated into a Nikon Eclipse Ti microscope through a C-mount. For QPM imaging, we used a halogen lamp as the transmitted light source. A Nikon LWD N.A. 0.52 condenser was used with the aperture diaphragm minimized. A C-HGFI mercury lamp was used for fluorescence illumination. A Nikon TI-S-ER motorized stage was used to position the sample with the moving speed of 2.5 mm/s in XY direction (accuracy 0.1 µm). A Nikon Perfect Focus System (PFS) was used for maintaining the focus. An Endow GFP/EGFP filter sets (Chroma 41017) was used to take the Geminin-GFP image. We used three objective lenses as indicated in this study, one Nikon Plan Flour 10X N.A. 0.3 PFS dry, one Nikon Plan Apo 20X N.A. 0.75 PFS dry, and one Nikon Plan Apo 40X N.A. 0.95 PFS dry. NIS-Elements AR ver. 4.13.0.1 software with the WellPlate plugin was used to acquire images. A homemade incubation chamber was used to maintain the constant environment of 36°C and 5% CO2.

### Quantification of QPM measurement errors

To quantify the OPD noise of the blank sample, we performed all the measurements as described on the blank 6-well glass-bottom plates filled by Phosphate-Buffered Saline (PBS, Corning, 21-040-CV) and covered with mineral oil (Sigma-Aldrich, M8410) at 10X magnification.

Fixed cells were used to quantify the dry mass and growth rate measurement error. For sample preparation, HeLa cells were seeded in 6-well glass-bottom plates (Cellvis, P06-1.5H-N) at 3500 cells/cm^2^ overnight, then fixed in 0.2% glutaraldehyde (50 wt. % in water, Sigma-Aldrich, 340855) for 10 min at room temperature. Then the fixed cells were immersed in PBS and topped with mineral oil.

In the experiments to quantify the dry mass measurement error, cells were imaged every 5 min for 2 hours. At 10X magnification, an area of 8X8 FOVs in the well center was scanned, with the X-Y step distance as one-fifth of the FOV. At 20X, an area of 15X15 FOVs was scanned, with one-fifth of the FOV as the step distance. At 40X, 60 cells were chosen manually; each was imaged in four FOVs with the cell at a different corner. The temporal error was quantified as the standard deviation of the time series of each cell divided by the mean mass of the cell. To quantify the net spatial error, we averaged the dry mass measurements through the time series first to eliminate the error caused by the temporal variation, then took the standard deviation divided by the mean of each cell at different positions as the spatial error. The temporal and spatial combined error was the standard deviation divided by the mean of each cell at different positions without averaging by time series.

### Long-term live-cell imaging under QPM

The 6-well glass-bottom plates were treated by Plasma Etcher 500-II (Technics West Inc.) at 75 mTorr, 110 W, for 1 min, then coated by 50 µg/mL fibronectin (Sigma-Aldrich, F1141) overnight. Cells were seeded on the pre-coated plates at 2000 cells/cm^2^ 3 hours prior to the experiments in the medium of DMEM without phenol red (ThermoFisher Scientific, 21063) supplemented with 10% FBS, 1% penicillin/streptomycin, and 10 mM sodium pyruvate, and topped with mineral oil. All the experiments were done at 10X magnification. The Phase images were acquired every 30 min, and the fluorescent images were acquired every 1 hour, at 36°C by the SID4BIO camera. The HeLa cells in normal growth condition or with thymidine were monitored for 40-48 hours. The RPE cells were monitored for 36-45 hours. The HeLa cells with Rapamycin were monitored for 70-72 hours. The drugs were added just before the time-lapse movie. The position of the FOVs was generated by a custom-developed GUI in Matlab (MathWorks), which assured that the FOVs were within the center 10% area of the well and were evenly spaced. 72 FOVs were imaged in each well. The fixed cells used to analyze instrumental fluctuation were imaged every 30 min for 27 hours.

### QPM Image analysis and data processing

We developed a custom GUI to evaluate the performance of the synthetic reference and parameters for segmentation. All the image processing pipeline (generating the reference wavefront, applying the reference, background subtraction, cell segmentation, and cell tracking) was conducted on a high performance compute cluster by custom-written codes in Matlab. To get the most accurate growth rate measurement, we cleaned up all the data points of contact or rounded cells. This data cleaning process created gaps in the cell tracking trajectories. We discarded all the trajectories with any single gap longer than 3 hours or total gap longer than 6 hours. The cleaning process also removed the data immediately after birth and before mitosis. Thus we considered any trajectory that begins less than 3 hours after birth and ends less than 1 hour before mitosis as a trajectory of the full cell cycle. The gaps in the trajectories were filled by linear interpolation. For HeLa cells arrested by thymidine, any trajectory longer than 12 hours and with an identified G1/S transition was adopted. For each condition, the trajectories were collected from more than three independent experiments except for the thymidine treatment, which was done only once.

To get the single-cell growth rate trajectory, we applied cubic spline smoothing (the csaps function in Matlab) on the dry mass trajectories at the smoothing parameter *p* = 0.00002 and then fitted the linear growth rate in a 3-hour sliding window to further reduce noise. The same processing was applied to fixed-cell data after levelling to demonstrate their effect on the power spectrum.

The dry mass trajectories were fitted by the linear, exponential, or second-order polynomial functions using linear (for the linear function) or nonlinear least squares (for the exponential and second-order polynomial functions) fit with bisquare weighting to eliminate the impact of outliers. The goodness of fit was estimated by the small sample Akaike Information Criterion AICc(44).

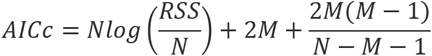

where *RSS* is the Residual Sum of Squares of the fit, *N* is the number of data points, and *M* is the number of parameters in the function.

The dry mass, growth rate, or detrended dry mass trajectories were aligned and averaged as indicated. When the trajectories were aligned to birth, G1/S, or chronological time, the last 2 hours before division was trimmed to avoid the impact of the abrupt mitotic dip. In the thymidine treated data, since the detrended dry mass trajectory was too noisy, the first peak after G1/S was identified by the smoothed growth rate trajectory. Since the trajectory length varied a lot among cells, the mean trajectory only included data points of the average of more than 50 trajectories except for the dataset of thymidine treatment, where the threshold was reduced to 25 due to the low cell number. The 95% confidence intervals of the mean trajectory were calculated as 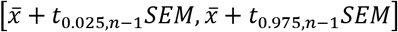 where n is the trajectory number; *SEM* is the Standard Error of the Mean,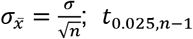 and *t*_0.975,*n*−1_ are the t-scores at 2.5% and 97.5% tails with degree of freedom equal to *n* − 1.

### SMR data analysis

The L1210 data were adopted from the L1210 FUCCI control dataset measured by the small-channel SMR in Mu et. al.(49). A total number of 63 cells measured in 9 independent experiments were analyzed. As the time interval of the SMR data was irregular with a mean at 1.1 minutes, we first linear interpolated the data at a fixed time interval of 1.2 minutes. The buoyant mass trajectories were fitted by the linear, exponential, and polynomial functions. The goodness of fit of different functions was compared by the Akaike Information Criterion(64),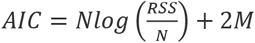. All the trajectories were detrended by the third-order polynomial in further analysis. The robust periodogram of the detrended trajectories was estimated and the periodicity was tested by the robust detection method. The detrended trajectories were fitted by the generic cosine function,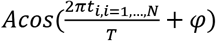, using the nonlinear least-square fit. The last 2 hours before division was trimmed before fitting to remove the mitotic dip. The frequency of the maximum peak in the periodogram was adopted to calculate the initial value of T for the fitting.

### Periodogram and robust periodicity detection

The robust periodogram was estimated by the correlogram as described in Ahdesmäki et al. (47). In the cases of QPM trajectories aligned to cell division, the last 1 hour before division was trimmed off from the mean trajectory to avoid the dramatic impact of the mitotic dip on the periodogram. The last 2 hours of the SMR trajectory aligned to mitotic dip were trimmed off for the same reason. For the thymidine treated cells aligned to the first peak after G1/S, the mean trajectory after the first dip after G1/S was used to estimate the periodogram of the arrested S phase. All the significances were assigned by the permutation method. The implementation was realized by the Matlab source code provided in Ahdesmäki et al. (47) with slight modification. All the results of the statistical tests were summarized in Table S2.

## Acknowledgments

We thank the Nikon Imaging Center at Harvard Medical School for their courtesy in sharing space and other resources. We thank the National Institute of General Medical Sciences for support (5RO1GM26875-42). All the image processing, data transfer, and data storage were conducted on the O2 Compute Cluster, supported by the Research Computing Group at Harvard Medical School. The plasma treatment was done at the HMS Microfluidics Core Facility. We thank Teemu P. Miettinen, Joon Ho Kang, and Scott R. Manalis for generously providing the L1210 data and inspiring discussions; Harri Lähdesmäki and Miika Ahdesmäki for the help on the robust detection method and providing the source code, Ahmed Rattani, Doaa Megahed, Hong Kang, Johan Paulsson, Mingjie Dai, Scott Gruver, Scott Rata, Wenzhe Ma, and Ying Lu for helpful discussions.

