## Supplementary Information for "Computationally Enhanced Quantitative Phase Microscopy Reveals Autonomous Oscillations in Mammalian Cell Growth"

**This PDF file includes:**

Supplementary text
Figures S1 to S3
Tables S1 to S2
Legends for Movies S1

**Other supplementary materials for this manuscript include the following:**

Movies S1

**Supplementary Information Text**

**The goodness of fit of the linear and exponential growth model**

Although the measurement error is positively correlated with cell dry mass (A), the dispersion of the exponential fit does not carry the strong positive correlation (B), meaning the fluctuations in the cell mass trajectory are not solely due to the measurement error. Thus, we used the nonlinear least-square fit to fit the exponential growth model. Since the linear and exponential growth models have the same number of parameters, the goodness of fit only depends on their Residual Sum of Squares (RSS). We chose the small sample Akaike Information Criterion (AICc) to compare the goodness of fit of the two models(1). Using other criteria, such as the Bayesian Information Criterion (BIC), does not affect the result.

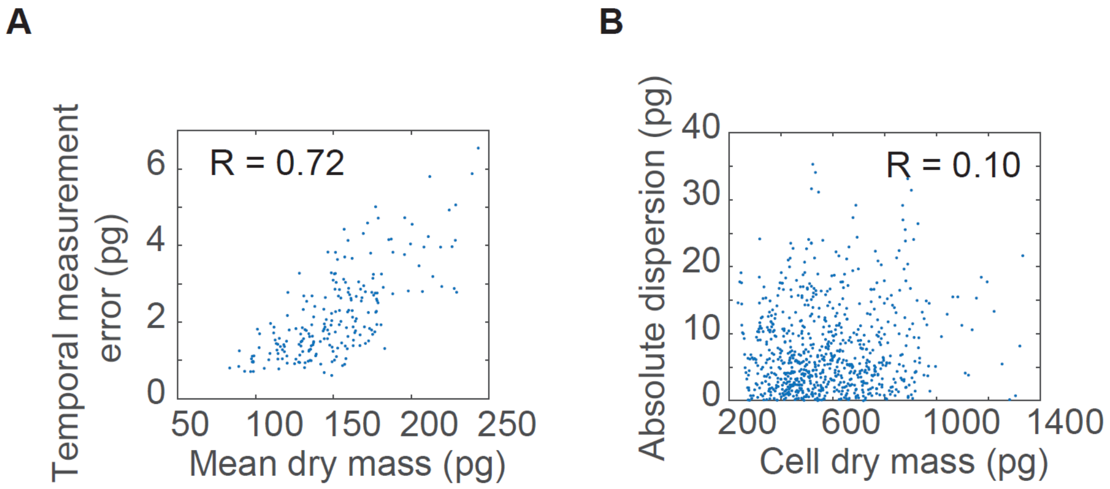


Figure. (A) The strong positive correlation between the mean dry mass and the temporal measurement error. Data is from Table 1 in the main text under 10X magnification. (B) The correlation between the cell dry mass and the absolute dispersion of the exponential fit. Data is from Fig. 4B in the main text with AICc_exp - AICc_li > 100; each data point is from one time point of a single-cell trajectory.

**The effect of data processing on the data power spectrum**

We used the long-term fixed-cell data to demonstrate how different data processing steps manipulate the power spectrum of the data and cause diminished or artificial information at certain time scales (see the figure below). The fixed cell trajectories were leveled by subtracting the mean and then underwent different data processing as indicated. The processed trajectories were stitched to estimate the Power Spectral Density (PSD) by using the Welch's method (Matlab's built-in function pwelch) with the segment length equal to 100. Taking the difference of adjacent points suppresses the low-frequency noise but greatly amplified the high-frequency noise (note the PSD is in log scale). Cubic spline as an imperfect low pass filter surpasses the high-frequency noise while preserves the low-frequency power. Linear fit in a sliding window or moving average suppresses noise at specific frequencies corresponding to the window size. Combining data smoothing and derivative taking creates an artifactual peak in the power spectrum.

Data processing, especially noise filtering, is commonly applied to single-cell dynamics. But its use is rarely thoroughly examined analytically. The live-cell data carries the information of regulatory changes, stochastic process, and measurement error in the cells over time as well as cell-to-cell variability, which makes it difficult to investigate the effect of data processing objectively. Thus, we recommend to perform the power spectral analysis routinely with a control dataset, such as the fixed-cell data, to dissect potential error and misinterpretation.

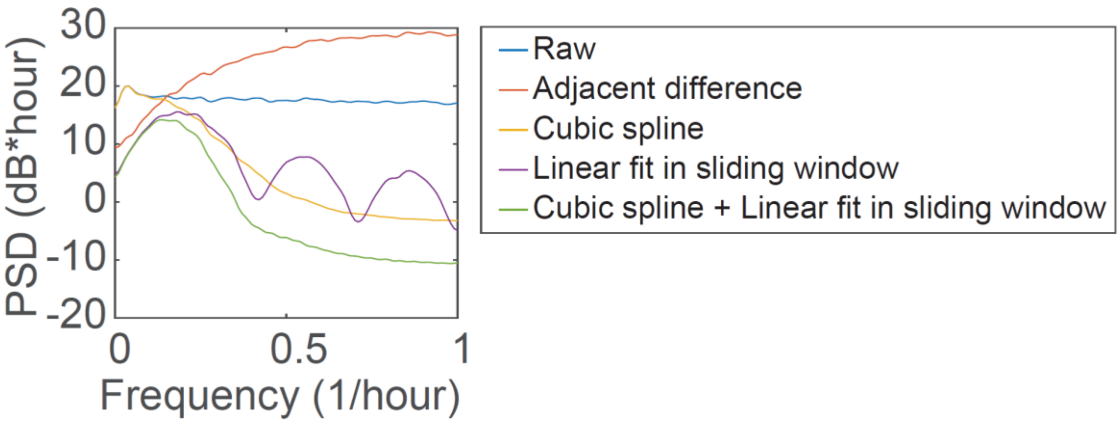


Figure. The effect of data processing on the power spectrum of fixed-cell data. The smoothing parameter of the cubic spline is 0.00002; the sliding window of the linear fit is 3 hours. The growth rate trajectories in Fig. 4C were calculated by the same processing steps as in the green spectrum.

**False Detection Rate (FDR) of g- and g'-test**

We tested the FDR of the conventional g-test and the modified g'-test with the random trajectories of Gaussian noise as well as the mean trajectories of randomly permutated single-cell trajectories. The g-test discovered 23 out of 500 cases (4.6%) in random trajectories as oscillatory with p-Value smaller than 0.05, while the g'-test with the truncated frequency window $f_l > 0.1/\mathrm{hour}$ discovered 29 out of 500 (5.8%). Similarly, the g'-test discovered 20 out of 500 oscillatory cases (4.0%) in the mean trajectories of permutated single-cell trajectories (using the HeLa dataset of 340 trajectories), compared with 21 out of 500 cases (4.2%) by the g-test. In summary, we concluded the FDR of the g- and g'-test were well calibrated.

**Detrend the SMR data**

We tried different fitting functions to detrend the SMR data and compared their goodness of fit by the Akaike Information Criterion (AIC)(2). The smaller AIC represents the better fit. The third order polynomial fits better than the linear or exponential models in all the trajectories, and the fourth order polynomial does not improve the fit significantly than the third order polynomial (see the figure below). Thus, we detrended the SMR trajectories by the third order polynomial fit.

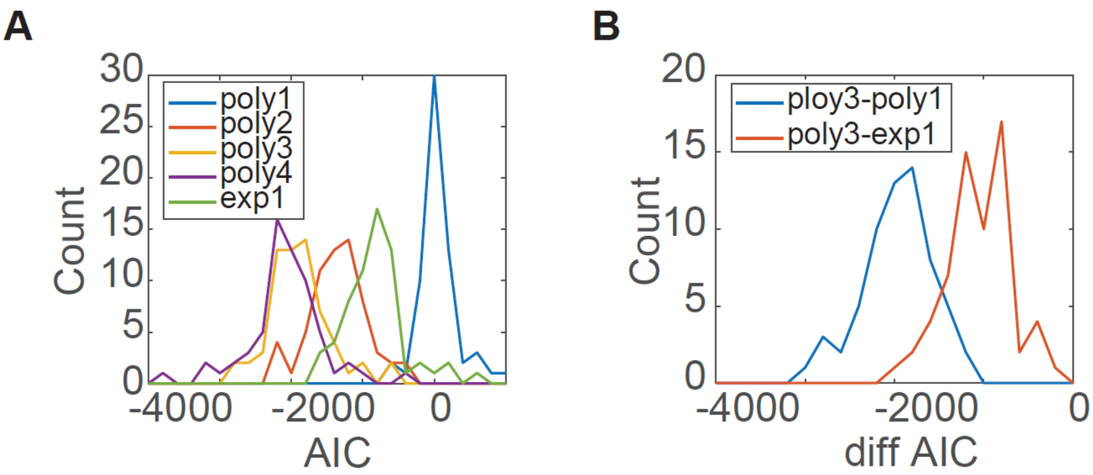


Figure. (A) The histogram of the AIC of different fitting functions. (B) The histogram of the AIC difference between the third order polynomial and the linear (blue) or exponential (red) model.

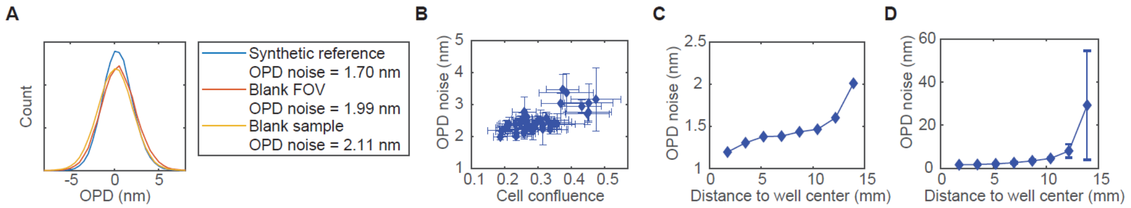


**Fig. S1.** (A) The OPD noise (the standard deviation of the OPD in the blank area) of an FOV on a blank sample when applying a reference from the synthetic method (blue), its neighbor FOV (red), or an FOV of the same light path on another blank sample (yellow). (B) The OPD noise changes with the reference made by cell FOVs of different confluence. Each data point is from a single experiment; the error bars indicate the standard deviations of all the FOVs measured in that experiment. (C) The OPD noise of a FOV at the center of a blank 35 mm well shows small increase when the synthetic references are made with FOVs at increasing distance to the well center. (D) When applying the reference made by the central 1% area of the well, the OPD noise of a blank 35 mm well is less than 3.7 nm up to 10 mm distance to the well center. Error bars indicate the standard deviations of the FOVs at a certain distance.

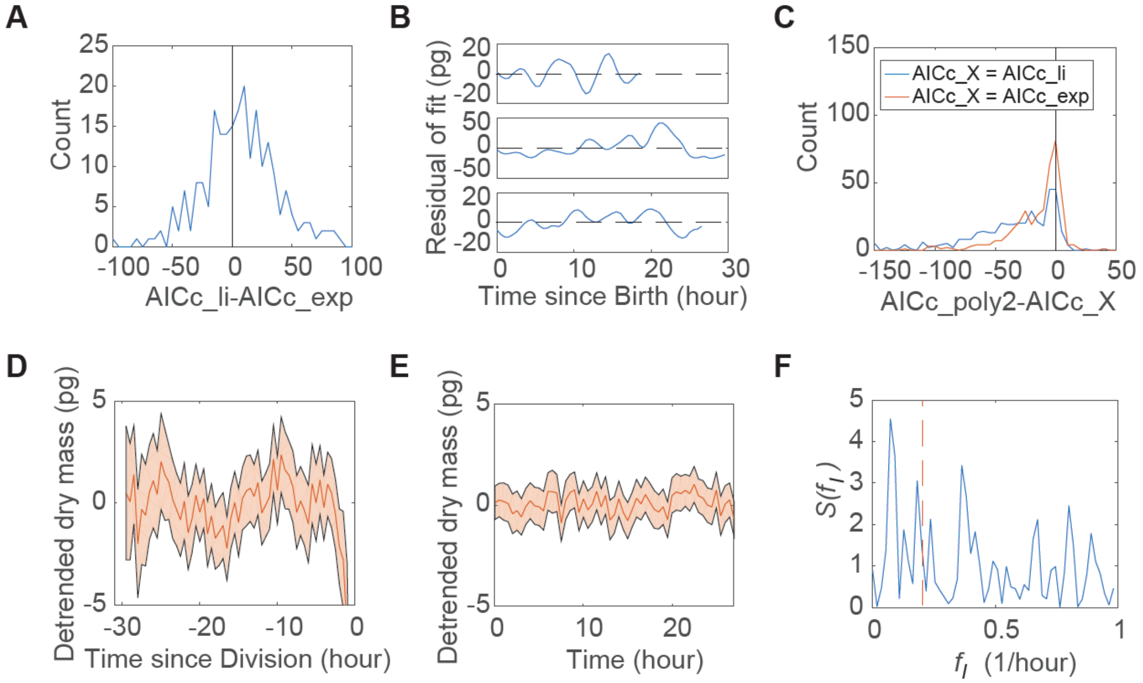


**Fig. S2.** (A) The histogram of the difference between the AICc of the linear (AICc_li) and exponential (AICc_exp) fit in RPE cells. (B) Examples of the residual trajectories, when the smoothed dry mass trajectroies of HeLa cells were fitted by the exponential model. The horizontal dashed lines indicate residual equal to zero. (C) The histogram of of the difference between the AICc of the second-order polynomial fit (AICc_poly2) and the linear or exponential fit (AICc_X, X=li or exp, respectively). The black vertical line indicate the difference equal to zero. (D) The mean behavior of detrended dry mass trajectories of HeLa cells aligned to cell division time. The red shaded region indicates the 95% confidence intervals of the mean. n = 340. (E) The mean behavior of detrended dry mass trajectories of fixed cells aligned to their chronological time. The red shaded region indicates the 95% confidence intervals of the mean. n = 564. (F) The periodogram of the mean trajectory in (E). The red dashed line indicates the location of f=0.2/hour.

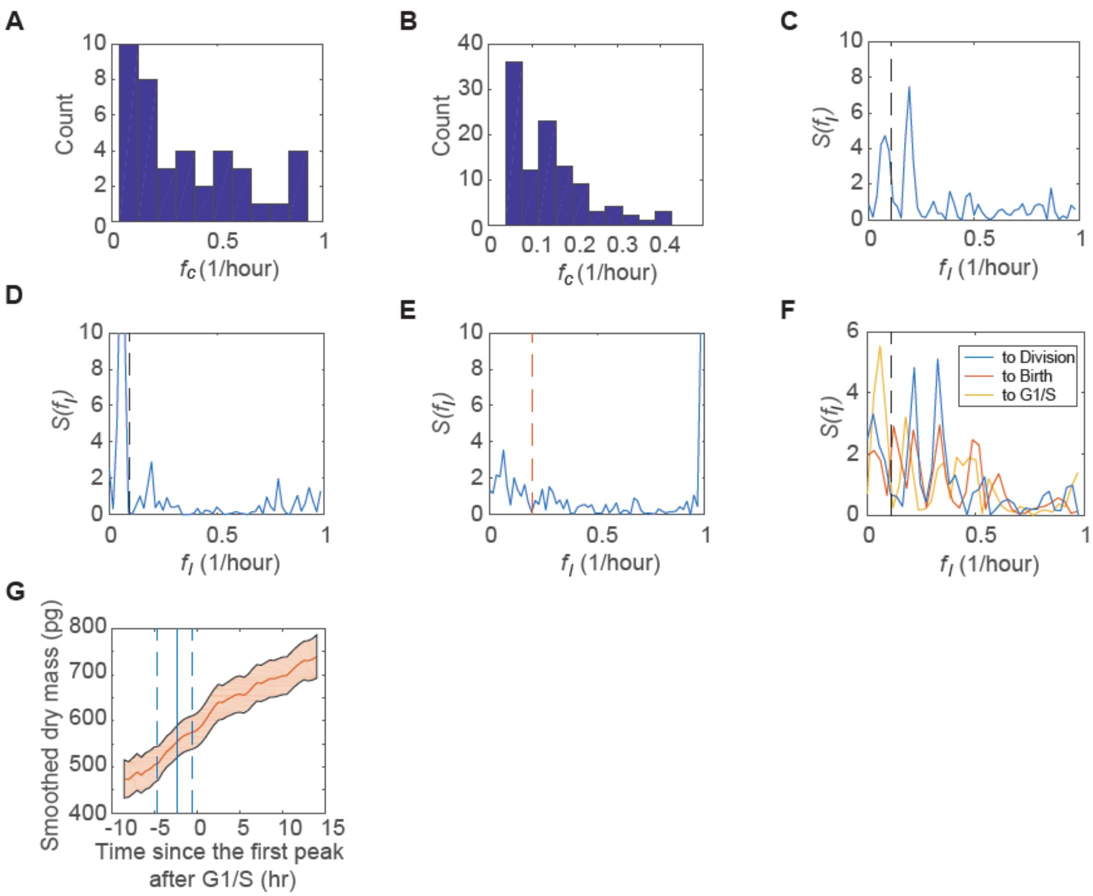


**Fig. S3.** (A, B) The histogram of the oscillation frequency, $f_c$, of the fixed cells (A) or live HeLa cells (B) that were identified to be oscillatory by the robust detection method. (C) The periodogram of the mean trajectory of HeLa cells aligned to their division time, after removing all the oscillatory individuals. n = 234. (D, E) The periodogram of the mean trajectory of HeLa cells from a single experiment aligned to cell division (D) or chronological time (E), respectively. n = 170. (F) The periodogram of the mean trajectory of RPE cells aligned to cell division (blue), birth (red), or G1/S (yellow), respectively. n = 197. (G) The mean trajectory of smoothed dry mass of HeLa cells under the treatment of thymidine aligned to the first peak after G1/S; n = 51. The vertical blue line indicates the averaged timing of G1/S; the dashed lines are the 5 and 95% percentile. The red shaded region indicates the 95% confidence intervals of the mean. The black dashed lines in (C, D, and F) indicate $f_{min}$=0.1/hour used in the g'-statistic test. The red dashed line in (E) indicates the location of $f$=0.2/hour.

Table S1. The weight parameters used to track HeLa cells.

| $w_D$ | $w_M$ | $w_A$ | $w_G$ |
|---|---|---|---|
| 0.03 | 2 | 0.05 | 0.1 |

Table S2. The summary of the statistical test results of the mean trajectories by the robust detection method. $p1 = p(g>g_o)$, $p2 = p(g'(f_l>0.1)>g'_o(f_l>0.1))$, $p3 = p(g(f_l=f_x)>g_o(f_l=f_x))$, where $f_x$ is the closest frequency to 0.2/hour in $f_l$.

| Experimental condition | Dataset | Alignment point | n | N | fc (1/hour) | Period (hour) | p1 | p2 | p3 | Average cell cycle length (hour) |
|---|---|---|---|---|---|---|---|---|---|---|
| HeLa, normal | All the trajectories | Division | 340 | 57 | 0.1930 | 5.18 | 0.0014 | <0.0002 | | 26.06 |
| | | Birth | | 54 | 0.1852 | 5.40 | | 0.0410 | | |
| | | G1/S | | 55 | | | | 0.2849 | | |
| | Without the oscillatory individuals | Division | 234 | 52 | 0.1923 | 5.20 | | 0.0018 | | |
| | From a single experiment | Division | 170 | 56 | 0.1964 | 5.09 | | 0.0632 | | |
| | | Chronological time | | 59 | 0.9831 | 1.02 | <0.0002 | | 0.9999 | |
| | Randomly selected | Division | 188 | 51 | 0.1961 | 5.10 | | 0.0016 | | |
| HeLa, fixed | All the trajectories | Unaligned | 564 | 55 | | | 0.4334 | | 0.9999 | |
| HeLa, 2mM thymidine | All the trajectories | First peak after G1/S | 51 | 26 | 0.1923 | 5.20 | | <0.0002 | | |
| | | Chronological time | | 43 | | | | 0.7622 | | |
| HeLa, 100nM Rapamycin | All the trajectories | Division | 188 | 70 | 0.2286 | 4.38 | | 0.2487 | | 34.97 |
| RPE, normal | All the trajectories | Division | 197 | 37 | 0.3243 | 3.08 | | 0.0794 | | 18.9 |
| | | Birth | | 33 | | | | 0.7856 | | |
| | | G1/S | | 34 | | | | 0.3647 | | |
| L1210, normal | All the trajectories | Birth | 63 | 419 | 0.2387 | 4.19 | <0.0002 | | | 10.42 |
| | | Mitotic dip | | 422 | 0.1777 | 5.63 | <0.0002 | | | |

**Movie S1 (separate file).** HeLa cells monitored by QPM for 48 hours. Each frame is a montage of 8 X 9 OPD images of 1.184 mm X 0.888 mm FOV observed under 10X objective. The time interval between frames is 30 minutes.
