## Supplementary figures and images for "Computationally Enhanced Quantitative Phase Microscopy Reveals Autonomous Oscillations in Mammalian Cell Growth"

### Movie S1

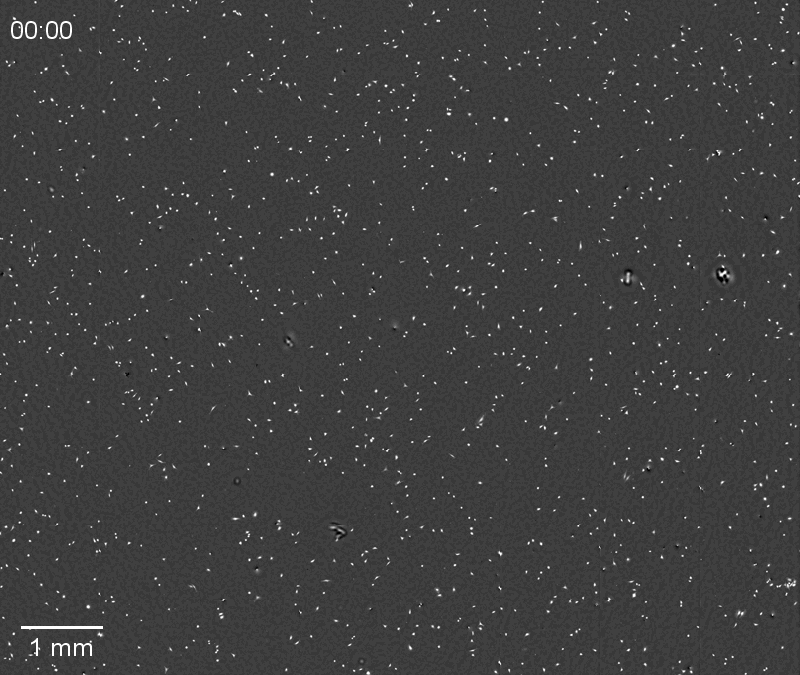
